# Monopolar orientation of kinetochores at meiosis is enforced by COHESINS and their regulators, CENP-C, and the deSUMOylase SPF2

**DOI:** 10.1101/2025.08.19.671082

**Authors:** Dipesh-Kumar Singh, Alexander Mahlandt, Sylvie Jolivet, Stephanie Durand, Birgit Walkemeier, Christelle Taochy, Victor Solier, Maria Derkacheva, Laurence Cromer, Raphael Mercier

## Abstract

The first division of meiosis is unique in its capacity to halve the ploidy of the future gametes. To this end, one key innovation compared to mitosis is the monopolar orientation of the pairs of sister kinetochores required for the proper separation of homologs at meiosis I. How monopolar orientation is imposed is unclear and seems to vary in eukaryotes. Here we performed a forward genetic screen in *Arabidopsis thaliana*, specifically designed to identify the molecular components imposing monopolar orientation, based on mutants’ ability to restore fertility in *spo11 osd1* haploid plants. We show that monopolar orientation involves all four cohesin subunits (REC8, SCC3, SMC1, SMC3), the cohesion establishment factors CTF18 and DCC1, and the cohesin protectors SGO1/2 and PANS1, the inner kinetochore protein CENP-C, and the desumoylase SPF2. The mutants show bipolar orientation of achiasmatic chromosomes, but monopolar orientation is maintained in the presence of crossovers despite splitting of sister kinetochores at metaphase I and reduced levels of cohesin. Taken together, the findings demonstrate that cohesion establishment and protection, kinetochore function, and deSUMOylation, together with crossovers, enforce monopolar orientation in plants and support a cohesion-driven model of kinetochore orientation at meiosis I that is conserved across kingdoms.

## Introduction

Meiosis is a cell division that promotes chromosomes to exchange genetic information and reduce their number by pairing homologous chromosomes and executing two rounds of division. Despite major strides in recent decades, our understanding of how exactly meiosis ensures replicated chromosomes are made to segregate together at one division and divide only at the next remains incomplete ^1,2^. A central player in the process is the kinetochore, a large protein complex that associates with centromeres and enables the attachment of microtubules to chromosomes. During meiosis I, the kinetochores of replicated sister chromatids act as a unified microtubule-binding face that ‘mono-orient’ towards one spindle pole, as opposed to ‘bi-orienting’ during mitosis. This process is known as monopolar orientation, or sister kinetochore co-orientation. The outcome of this co-orientation after the onset of the first division is the segregation of homologous chromosomes to opposing poles rather than chromatids, representing a reductional rather than equational division that allows for the later separation of sisters in pursuit of haploid spores. Essential for chromosome segregation are the activities of the Cohesin complex, a member of the SMC complex family, ring-shaped complexes that are involved in the packaging and organization of chromosomes during mitosis and meiosis ^3^. Cohesin is responsible for creating cohesion between replicated sister chromatids during S phase ^4^. During meiosis, a population of cohesin contains a meiosis-specific ‘kleisin’ subunit Rec8, and together with non-REC8-containing cohesin entraps sister chromatids. Rec8-containing cohesin in the vicinity of peri-centromeres is protected from removal by the protease Separase during anaphase I, which is crucial for maintaining sister chromatid cohesion until anaphase II and their balanced segregation ^5^. A conserved family of proteins called Shugoshins drives this protection by recruiting the phosphatase PP2A to peri-centromeric Rec8 ^6–9^. While clear that this protected cohesin is necessary to maintain sister chromatid cohesion after the first division, the absence of Rec8 also abolishes sister kinetochore co-orientation in fission yeast, plants, and vertebrates ^10–13^. A common mechanism has been proposed that suggests differences in cohesion between sister chromatids along their arms and at their core- and peri-centromeres during meiosis are fundamental to establishing the monopolar orientation of sister kinetochores before the first meiotic division ^1,14–17^. This presents a model in which Rec8 at the core centromere but not the peri-centromeres is crucial for the co-orientation of sister kinetochores, supported by findings that the depletion of core-centromeric Rec8 causes bi-orientation in mouse^18^. Aurora kinase is involved in correcting microtubule-kinetochore attachments during mitosis, and similarly works to promote correct attachments to mono-orienting kinetochores during meiosis I ^19^. The role of cohesin is not, however, unique, as point-centromere containing budding yeast harbors a protein complex called Monopolin containing a meiosis-specific subunit Mam1, required for monopolar orientation in *S. cerevisiae* ^20^. Monopolin directly cross-links sister kinetochores and drives co-orientation independent of cohesion; functional homologs of Mam1 have not been identified outside of *S. cerevisae* ^20–23^.

Notably, meiosis-specific proteins that localize to kinetochores have been identified in yeast and vertebrates that drive co-orientation in a cohesion-dependent manner; mouse Meikin, fission yeast Moa1, and budding yeast Spo13 function in localizing Rec8-phosphorylating kinases directly to the kinetochore to promote core centromere cohesion ^13,16,24,25^. Functional homologs of Meikin/Moa1 nor any additional factors beyond the cohesin subunits Rec8 and SCC3 that have a role in co-orientation have not been identified in plants, raising the question of whether such a mechanism is broadly conserved.

Here, a genetic screening was undertaken in Arabidopsis to identify factors involved in monopolar orientation. Mutant alleles identified in Cohesin subunits, the Cohesin regulators CTF18-DCC1 and SGO1/SGO2, the kinetochore protein CENP-C, and the SPF2 deSUMOylase disturb co-orientation of kinetochores. Weakened monopolar orientation is associated with physical separation of sister kinetochores, reduced levels of chromosomal cohesins and premature loss of cohesion at meiosis II. These findings support a model in which centromeric Cohesins play parallel roles in monopolar orientation at meiosis I and sister chromatid cohesion at meiosis II.

## Results

### A forward genetic screen for monopolar orientation

We designed a specific genetic screen to identify factors promoting the monopolar orientation of kinetochores at meiosis I, based on the restoration of the fertility of haploid plants (Figure 1 and 2). In haploid Arabidopsis plants, meiosis still occurs but chromosomes cannot recombine because of the absence of homologs (Figure 1D) ^26,27^. At anaphase I, the resulting five univalents do not separate into chromatids like at mitosis (i.e., 5:5) but rather segregate as single units at anaphase I (e.g., 3:2), demonstrating that the monopolar orientation of the sister chromatid kinetochores is maintained in haploid meiosis. The resulting unbalanced first division is followed by a second division that distributes sister chromatids (Figure 1D). As a consequence, haploid plants have extremely low fertility. However, turning meiosis into mitosis in haploids with the *MiMe* mutations bypasses the meiotic defects and restores fertility (Figure 1E, ^27^). *MiMe* is composed of three mutations - *spo11, rec8* and *osd1*-that abolish the three key differences between meiosis and mitosis: recombination, the monopolar orientation of kinetochores, and the occurrence of the second division, respectively (Figure 1) ^28^. Of particular interest here, the double mutant *spo11 osd1* produced unbalanced spores and is sterile, in both haploids and diploids (Figure 1C, 1F), because the monopolar orientation of kinetochores prevents balanced segregation of sister chromatids. We thus performed a forward genetic screen in haploid *spo11 osd1* mutants with the rationale that any mutation that would affect monopolar orientation of kinetochores would restore fertility, as the *rec8* mutation does. Fertility is easy to assess visually by simply looking at the size of the fruits, making a large forward genetic screen feasible.

**Figure 1.**
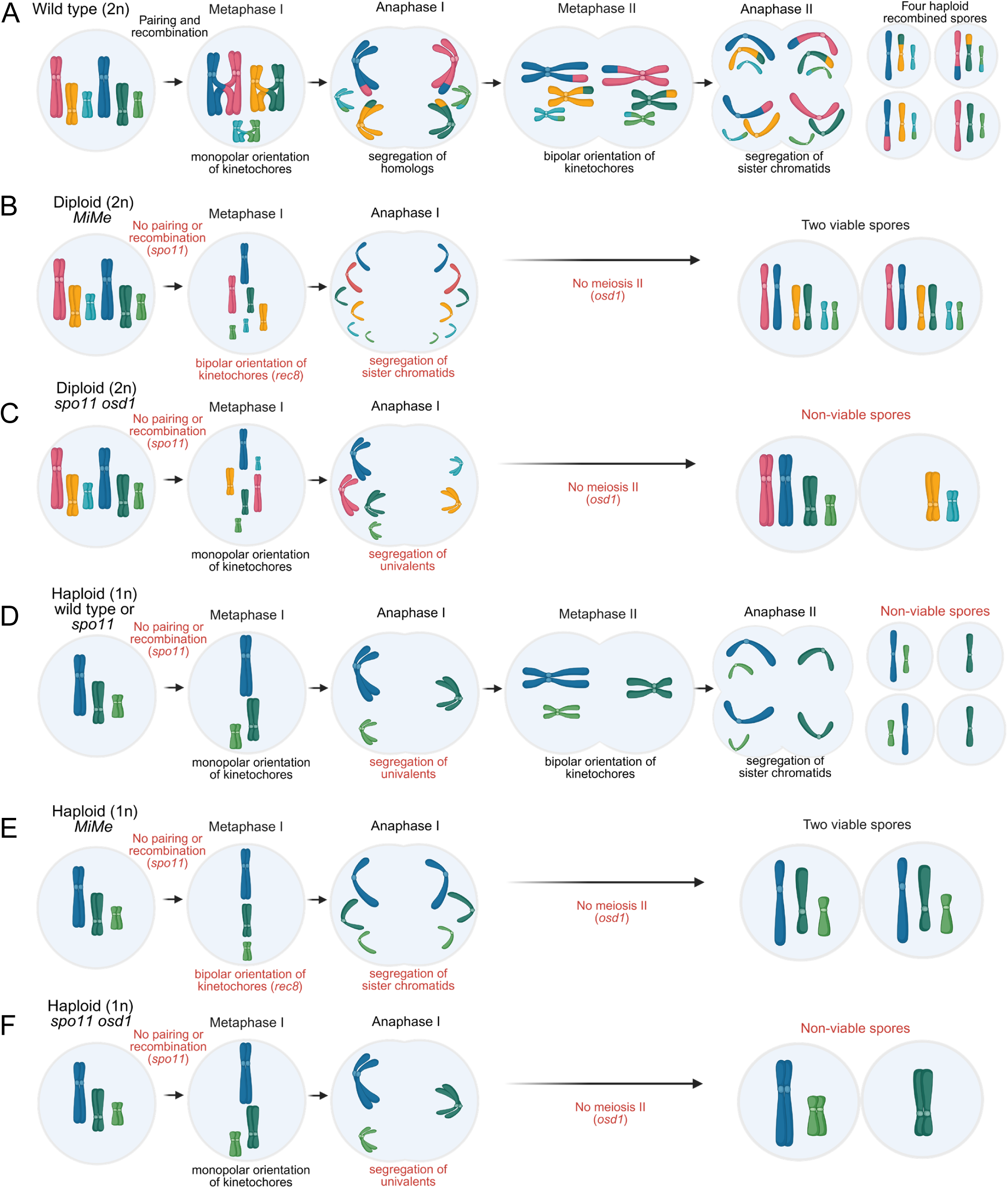
*MiMe* meiosis produces viable spores in diploid and haploid plants. Schematic description of meiosis in (A) the diploid wild type; homologous chromosomes undergo pairing and recombination, followed by two successive rounds of division to yield haploid recombined spores. (B) diploid *MiMe;* pairing, recombination, monopolar orientation, and the second division are lost due to mutations in *spo11, rec8,* and *osd1*, yielding unreduced and non-recombined spores. (C) diploid *spo11 osd1*; pairing, recombination, and the second division are lost, yielding non-viable spores that lack a full chromosome complement. (D), haploid wildtype or *spo11;* pairing and recombination are absent due to missing homologous chromosomes or double strand break formation, leading to unbalanced segregation and non-viable spores. (E) haploid *MiMe* (*spo11 osd1 rec8*); a full chromosome complement allows unreduced and non-recombined spores to be viable. (F) haploid *spo11 osd1;* pairing, recombination, and the second division are lost, yielding non-viable spores that lack a full chromosome complement. Note that three chromosomes are represented while A. thaliana has five.

**Figure 2.**
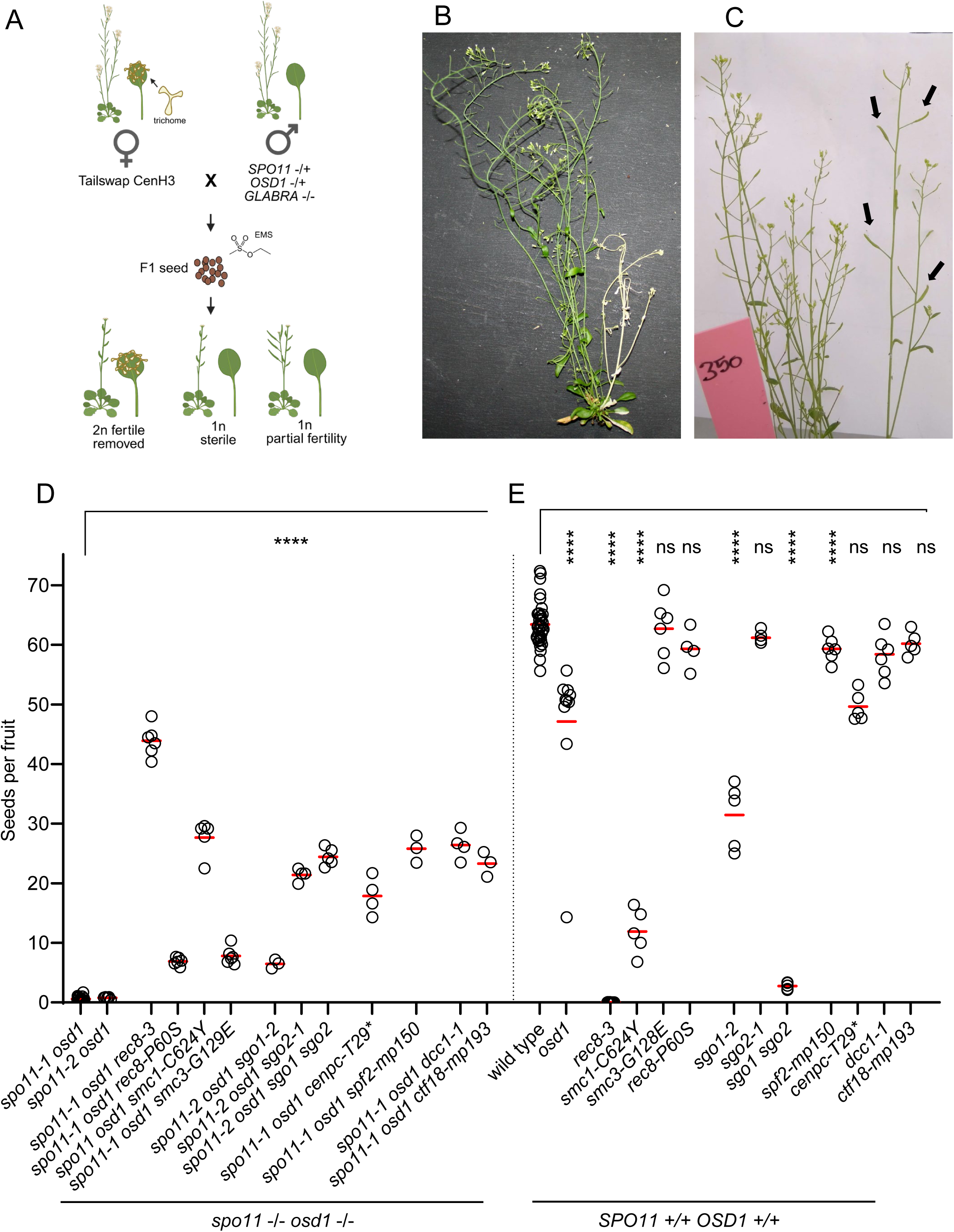
A forward genetic screen for monopolar orientation. (A) Principle of the genetic screen. (B) An M1 individuals displaying chlorosis in single branch. (C) A M1 mutant showing restored fertility (mp350). Arrows points to elongated fruits in a single branch, while the rest of the plants has short fruits. (D) Fertility of the identified monopolar orientation mutants in the *spo11 osd1* diploid contexts. Each dot represents the fertility of one plant, measured as the number of seeds per fruit counted on 10 fruits. The read line indicates the mean. (E) Fertility of the identified monopolar orientation mutants in the *spo11 osd1* diploid contexts. Legend as in D. Test are one-way ANOVA with Dunnet correction for multiple testing. ns: p>0.05. ****: p<10^-4^

In practice (Figure 2A), we took advantage of the TailSwap haploid inducer line ^29^, whose genome is eliminated in a proportion of embryos following crosses, generating haploid progeny containing only the genome of the other parent. We crossed the haploid inducer as female with plants heterozygous for *spo11-1* and *osd1* mutations and homozygous for the *glabra1* mutation, which confers a glabrous appearance to the leaves (Figure 2A). We applied moderate EMS mutagenesis on the seeds and selected the ∼30% of haploid plants based on the *glabra1* phenotype. Plants with trichomes were eliminated at an early stage of development, as the presence of the *GLABRA1* allele indicates normal fertilization and thus diploidy. One additional advantage of the screen in a haploid context is the phenotypic expression of recessive mutations in the first generation. As EMS mutagenesis is applied on seeds that contain several pluripotent cells, the resulting plants are chimeric for the induced mutations, as exemplified by chlorotic mutations observed in sectors of some M1 plants (Figure 2B). We screened ∼9000 *glabra* mutagenized plants, looking for branches with longer fruits, as candidates for carrying a mutation affecting the monopolar orientation of kinetochores (Figure 2C). Branches with longer fruits were genotyped for *osd1* and *spo11* and ploidy was tested through chromosome spreads on terminal buds. In many cases, these were diploid, despite having the *glabra* phenotype. This noise could result either from the spontaneous doubling of somatic cells ^26^, or more complex genetic events associated with genome elimination ^30^. In total, we retained thirteen lines derived from haploid branches with enhanced fertility. A combination of whole genome sequencing, genetic mapping, and candidate testing with independent alleles led to the identification of the causal mutations in these thirteen lines, corresponding to ten genes (Table 1).

**Table 1.** Mutations recovered from the “monopolin” forward genetic screen.

| protein | function | allele | gene | position<br>(TAIR10) |  | effect on the<br>protein | method of<br>confirmation | comment |
| --- | --- | --- | --- | --- | --- | --- | --- | --- |
| REC8 | Cohesin<br>subunit | mp257 | AT5G05490 | Chr5:1624739 | C>T | R20>STOP | independent<br>alleles | meiosis specific |
|  |  | mp305 |  | Chr5:1625214 | C>T | P60>S |  |  |
| SMC1 | Cohesin<br>subunit | mp172 | AT3G54670 | Chr3:20239572 | G>A | C624>Y | complementation<br>test | null mutation is<br>lethal |
| SMC3 | Cohesin<br>subunit | mp226 | AT2G27170 | Chr2:11615684 | C>T | G129>E | mapping to a<br>single mutation | null mutation is<br>lethal |
| SCC3 | Cohesin<br>subunit | mp225 | AT2G47980 | Chr2:19633111 | A>G | R326>G | independent<br>alleles | null mutation is<br>lethal |
| CTF18 | cohesin<br>loading | mp193 | AT1G04730 | Chr1:1327905 | C>T | splice site | independent<br>alleles |  |
|  |  | mp274 |  | Chr1:1326791 | C>T | splice site |  |  |
| DCC1 | cohesin<br>loading | mp270 | AT2G44580 | Chr2:18403182 | C>T | splice site | independent<br>alleles |  |
| SGO2 | cohesin<br>protection | mp267 | AT5G04320 | Chr5:1210471 | G>A | splice site | independent<br>alleles |  |
| SGO1 | cohesin<br>protection | mp350 | AT3G10440 | Chr3:3248542 | G>A | splice site | independent<br>alleles |  |
| CENP-C | kinetochore | mp221 | AT1G15660 | Chr1:5381313 | ins C | T29 frameshift,<br>ATG at codon 56 | mapping to a<br>single mutation | null mutation is<br>lethal |
| SPF2 | desumoylase | mp150 | AT4G33620 | Chr4:16150537 | del C | L385 frameshift | independent<br>alleles |  |
|  |  | mp268 |  | Chr4:16150107 | G>A | D303>N |  |  |

### Quantification of monopolar orientation

We tested and quantified the effect of the identified mutations on monopolar orientation, through (i) their ability to restore fertility in diploid *spo11 osd1* plants (Figure 2D). Indeed, a monopolar mutation restoring fertility of haploid *spo11 osd1* is also expected to restore fertility of diploid *spo11 osd1* (Figure 1B). Note that the haploid *MiMe*-like plants found in the screen produce haploid gametes, and thus spontaneous diploid progeny. (ii) Second, we analysed the ability of the mutations to modify chromosome segregation at anaphase I in diploid *spo11* (i.e; not mutant for *osd1,* allowing observation of metaphase II. Figure 3). In diploid *spo11-1 or spo11-2,* the absence of recombination leads to the presence of ten univalents at metaphase I (figure 3D), which because of monopolar orientation, mostly segregate as single units at anaphase I, without separation of the sister chromatids, resulting in the erratic distribution of the ten univalents (e.g. 7:3) (Figure 1C, Figure 3D-F) ^31^. Occasionally, we observed the separation of one or two pairs of sister chromatids, indicating that univalents can infrequently orient in a bipolar manner at metaphase I in *spo11-1 or spo11-2*, and resulting in “mixed segregation” (Figure 4). This unbalanced segregation at meiosis I causes a quasi-sterility of the *spo11-1* and *spo11-2* mutants, alone or in combination with *osd1,* that skip the second division (Figure 2D). A mutation that abolishes monopolar orientation is expected to provoke the biorientation of univalents and distribution of sisters in *spo11*, and thus to restore fertility of diploid *spo11 osd1* mutants. We confirmed these predictions in the mutations identified in the genetic screen (see below).

**Figure 3.**
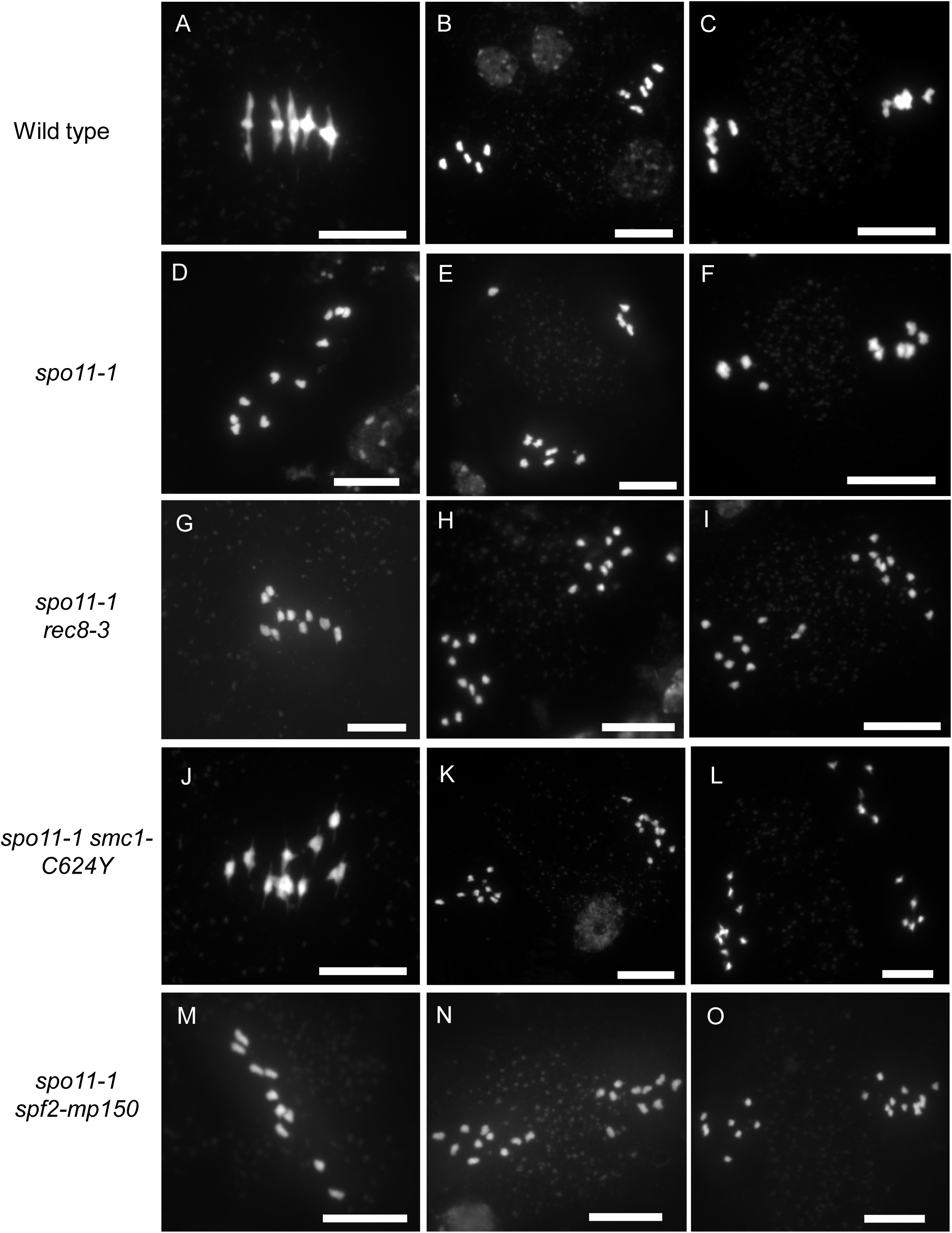
Mutation in *rec8*, smc1 or *spf2* leads to the bipolar orientation of univalents and distribution of sister chromatids. (A-I) Chromosome spreads of male meiocyte. (A) Wild-type metaphase I, with five bivalents aligned on the metaphase plate. (B) Wild-type metaphase II. Five chromosomes (pairs of chromatids) are aligned on the two metaphase II plates. (C) Wild-type anaphase II, with balanced distribution of the chromatids. (D) *spo11* metaphase I. Ten univalents are scattered. (E-F) *spo11-1* metaphase II with unbalanced segregation of univalents (7:3). (G) *rec8-3 spo11-1* metaphase I. Univalents align at metaphase I, suggesting bipolar orientation. (H-I) *rec8-3 spo11-1* metaphase II. (J) *smc1-C624Y spo11-1* metaphase I. (K-L*) smc1-C624Y spo11-1* metaphase II. (M) *spf2-mp150 spo11-1* metaphase I. (N-O) *spf2-mp150 spo11-1* metaphase II. Scale bar: 10 µm.

**Figure 4.**
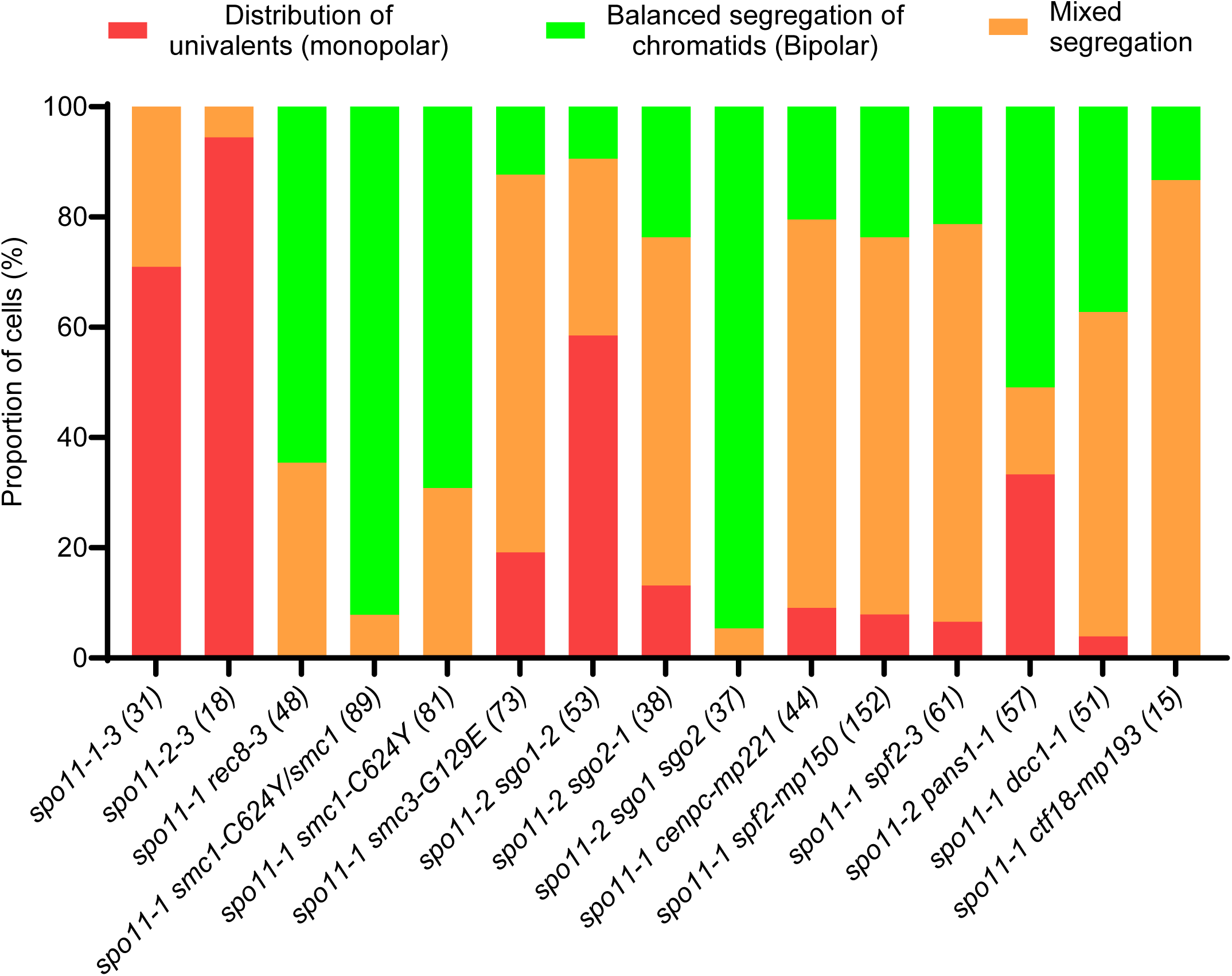
Quantification of chromosome segregation patterns. Proportion of cells displaying either monopolar, bipolar, or mixed segregation. Number of cells analyzed are in parentheses.

### The four subunits of the Cohesin complex promote monopolar orientation

The cohesin subunits *REC8* and SCC3 were previously shown to be involved in monopolar orientation of kinetochores in Arabidopsis ^11^. Consistently, the screen described above identified two mutations in *REC8*, one introducing a STOP codon at position 20 and one with introducing a missense at codon 60 (P60>S) (Table 1). Another line contained a missense mutation in *SCC3* (R326>G). This confirms the efficiency of the screen and the importance of these two cohesin subunits in imposing monopolar orientation at meiosis I. In addition, causal mutations were found in the two other cohesin subunits, SMC1 and SMC3 (*smc1-C624Y*, *smc3-G129E,* Table 1). The causality of the *smc3*-*G129E* mutation was confirmed by genetic mapping, being the sole EMS-induced mutation in *mp226* that co-segregates with fertility of *spo11 osd1*. The causality of the *smc1-C624Y* mutation was confirmed through allelic testing with the previously characterized *smc1* null mutation *SALK_017437* ^32^. SMC1-C624Y falls within the conserved hinge domain required for cohesin’s stable association with chromosomes ^33^. As SCC3, SMC1, and SMC3 are each encoded by a single gene and essential for plant development ^11,32,34^, the mutations identified here must represent partial or separation-of-function alleles affecting meiosis without impairing plant viability.

The *spo11 osd1* double mutant has very low fertility, yielding less than one seed per fruit on average (Figure 2D). In contrast, the *spo11 osd1 rec8-3* mutant shows a high level of fertility with 44 seeds per fruit, similar to the single *osd1* mutant (t-test p=0.53). The *smc1-C624Y* mutation also increased the fertility of *spo11 osd1*, but to a lesser extent than *rec8-3* (Figure 2D, 28 seeds). The *rec8-P60S* and *smc3-G129E* mutations had a milder, but significant, effect on *spo11 osd1* fertility (Figure 2D). This suggests different penetrance in monopolar orientation, with *rec8-3* having the strongest effect, *smc1-C624Y* being intermediate*, and rec8-P60S* and *smc3-G129E* having the mildest effect.

We then observed chromosome behavior on male meiotic chromosome spreads. In *spo11-1*, the ten univalents fail to align on the metaphase I plate and mostly segregate at anaphase I as a single unit in an erratic manner (e.g. 7:3). Occasionally, a univalent splits into chromatids at anaphase I, resulting in a “mixed segregation” rather than pure “distribution of univalents” (Figure 3 D-F, Figure 4). In contrast, when *spo11-1* was combined with *rec8-3*, *smc1-C624Y* or *smc3*-G129E, we observed metaphase I with aligned univalents followed by balanced segregation of sister chromatids, resulting in a 10:10 segregation (Figure 3G-H, 3J-K, Figure S1, Figure 4). This modification of chromosome segregation compared to *spo11-1* was almost fully penetrant in *spo11-1 rec8-3 and spo11-1 smc1-C624Y,* with a large majority of the cells showing a 10:10 segregation, a minority showing a mixed segregation, and none showing a *spo11*-like distribution of univalents. In *spo11-1 smc3*-*G129E* (Figure 4), a similar modification of segregation pattern was observed, but with a larger proportion of mixed segregation (Figure S1, Figure 4), which is consistent with the lower restoration of fertility observed in *spo11-1 osd1 smc3*-*G129E* (Figure 2D). Altogether, this shows that all core Cohesin subunits promote the monopolar orientation of sister kinetochores at meiosis I. The central role of REC8 suggests that Cohesin complexes containing alternative α-Kleisins subunit (SYN2-4, ^32^) cannot sustain monopolar orientation at meiosis. Note that alternative α-Kleisins cohesin complexes must be active in *spo11 rec8* background to ensure sister chromatid cohesion at metaphase I, but do not impose monopolar orientation. SMC1, SMC3, and SCC3, which are single-copy and essential Cohesin subunits, may be more crucial than suggested by the effects of the inevitable partial alleles analyzed here. The *smc1-C624Y* mutation, which has a strong effect on monopolar orientation, also affects slightly plant growth. Altogether, this strongly reinforces the conclusion that the entire cohesin complex is crucial in imposing monopolar orientation at meiosis I in plants.

### SHUGOSHINs promote monopolar orientation, with SGO2 playing a dominant role

In *Arabidopsis*, two SHUGOSHIN homologs, SGO1 and SGO2, were shown to protect centromeric cohesion at meiotic anaphase I, which is required for proper segregation of chromatids at meiosis II ^7,8,35^. SGO1 and SGO2 protect centromeric cohesion partially redundantly, with SGO1 having a more important role than SGO2. In the screen described here, mutations affecting a splicing site in both *SGO1* and *SGO2* were identified (Table 1). The role of SGO1 and SGO2 in monopolar orientation was confirmed by introducing the independent T-DNA alleles *sgo1-2* and *sgo2-1* ^8^ into *spo11-2 osd1.* Interestingly, *sgo2* provoked a stronger restoration of fertility than *sgo1* (t-test p=5.10^-6^, Figure 2D). Consistently, chromosome spreads of male meiocytes in *spo11-2 sgo1* and *spo11-2 sgo2* showed that *sgo2* provoked a stronger reduction of monopolar orientation (Chi-square, p=7.10^-5^. Figure 4). This shows that SGO2 has a more crucial role than SGO1 in imposing monopolar orientation, while it is the opposite for centromeric cohesion protection. Combining *sgo1* and *sgo2* mutations further restored the fertility of *spo11-2 osd1* (p=0.011), and bipolar orientation (p<10^-6^) (Figures 2 and 4). Thus, both SGO1 and SGO2 promote monopolar orientation in a partially redundant manner, with SGO2 playing a more important role. The observation that SGO1 and SGO2 have opposite relative importance for monopolar orientation and cohesin protection advocates that these two proteins have distinct functions and could suggest that different pools of cohesins promote monopolar orientation and centromeric cohesion.

As cohesins and cohesin protectors appear to promote monopolar orientation, we reasoned that PATRONUS, an Arabidopsis Securin homolog which protects cohesins by antagonizing Separase ^8,36^, could also promote monopolar orientation. Indeed, the *pans1* mutation was able to promote the bipolar distribution of sister chromatids in *spo11-1* (Figure 4). Note that the *pans1* mutation cannot be found in our forward screen as *pans1* is synthetic lethal in combination with *osd1* ^37^.

### Cohesion establishment factors CTF18 and DCC1 promote monopolar orientation

The forward screen for monopolar function yielded two additional genes that interact with cohesin: *CTF18* and *DCC1*. These proteins are known to function together as part of the RFC^Ctf18^ complex to facilitate the establishment of sister chromatid cohesion during replication ^38–40^. Two alleles were isolated for *CTF18* and one for *DCC1*, all affecting splicing sites and thus presumably strong or null alleles (Table 1). A CRISPR allele was generated in DCC1 for independent confirmation, *dcc1-2*, which carries a 1163-bp deletion in the coding sequence and encodes only the first 29 of 388 amino acids. The *dcc1-2* and *ctf18-mp193* mutations restored fertility of *spo11-1 osd1* to intermediate levels, similar to *smc1-C624Y* or *sgo2-1*, but less than *rec8-3* (Figure 2D). Next, we observed male meiotic behavior using chromosome spreads. In *spo11*, univalents are randomly distributed due to monopolar orientation or occasionally show a mixed segregation at meiosis I (Figure 3-4). In *spo11-1 dcc1-1* and *spo11-1 ctf18,* segregation patterns corresponded to a clear reduction of monopolar orientation, with a proportion of cells showing balanced segregation of chromatids and the rest having a mixed segregation (Figure 4, Figure S1). This implicates both CTF18 and DCC1 in imposing monopolar orientation, likely *via* facilitation of cohesion establishment, although at an importance below that of the Cohesin meiotic subunit REC8.

### The inner kinetochore protein CENP-C promotes monopolar orientation and cohesion protection

Among the mutants recovered in the screen, one had an insertion of one base pair in the coding sequence of the kinetochore protein CENP-C. Genetic mapping to a single mutation confirmed that this mutation was causal in restoring fertility of *spo11-1 osd1* (Figure 2D). The *cenpc*-*T29\** mutation induces a frameshift at codon T29, encoding then a histidine residue immediately followed by a STOP codon. Intriguingly, the *cenpc*-*T29\** mutant is viable, while the CENP-C protein is presumably essential in *Arabidopsis* as in other eukaryotes ^41^. However, an ATG codon at position 56 may serve as an alternative translation start codon that would generate a CENP-C protein truncated of 55 amino-acids at its N terminus. The *cenpc-T29\** mutation restores fertility of *spo11-1 osd1*, at levels less than *rec8-3* but greater than *smc3-G129E* or *sgo1-2* (Figure 2D). We next looked at male meiosis by chromosome spreads. The *cenpc-T29\** allele is able to provoke a 10:10 balanced segregation of sister chromatids in the *spo11-1* mutant and show a high level of mixed segregation (Figure 4). The frequency of biorientation is thus clearly increased, but less than with *rec8-3* or *smc1-C624Y* (Figure 4, Figure S1). This is consistent with the intermediate restoration of fertility in *cenpc-T29* spo11-1 osd1* (Figure 2D). These findings demonstrate that the kinetochore itself plays a role in monopolar orientation and suggest that the N-ter of CENP-C is implicated.

### The deSUMOylase SPF2 promotes monopolar orientation

Finally, we identified two independent mutations in the desumoylase *SPF2* (At4G33620), a frameshift mutation at position 385 (*mp150*) and a missense at codon 303 (D303>N) (*mp268*). The At4G33620 gene encodes for one of the two Arabidopsis homologs of the yeast deSUMOylase ULP2 ^42^. It is known as SPF2 (SUMO PROTEASE RELATED TO FERTILITY 2), but was also named in the literature ULP2-like1 and ASP2 (Arabidopsis SUMO protease 2) ^42–44^. The *spf2-mp150* mutation provokes a frameshift upstream of the protease domain of SPF2, presumably abolishing its function. This allele significantly restored the fertility of *spo11-1 osd1* to levels similar to that obtained with *smc1-C624Y, sgo2, ctf18 or dcc1*, yet less than obtained with *rec8 (*Figure 2D). During male meiosis in *spo11-1* mutants, univalents randomly segregate during meiosis I. Introducing the *spf2-mp150* or the *spf2-3* mutation in *spo11-1* provokes equational 10:10 segregation of chromatids or mixed segregation in most of the cells (Figure 3-4). This corresponds to an intermediate penetrance of the bipolar orientation induction, similar to other mutants but less than *rec8* and concordant with the intermediate restoration of fertility of *spf2-mp150 spo11 osd1*. This result implicates the deSUMOylase SPF2 in monopolar orientation, suggesting that the deSUMOylation of one or several targets by SPF2 favors monopolar orientation.

### Chiasmata promote monopolar orientation

Next, we examined the phenotype of the mutant identified in the screen in the presence of active SPO11 and OSD1, i.e., in single mutants. REC8 is required for efficient double-strand break repair at meiosis, leading to chromosome fragmentation and full sterility of the single *rec8* null mutant (i.e. in the presence of DNA double-strand breaks generated by SPO11-1/SPO11-2) ^11,45^.

In contrast, rec*8-P60S, smc3*-G129E, *spf2-mp150*, *cenpc-T29\**, *sgo2-1, ctf18-mp193*, and *dcc1-1* single mutants undergo largely normal meiosis, with segregation of homologues at the first division and sister chromatids at the second division (Figure 5, Figure S2)^8^. Each of these mutants also show complete fertility with the exception of *cenpc-T29\**, which has a fertility reduction of 18% relative to the wild type despite apparent normal meiosis (Figure 2E, Figure 5). In these mutants, five bivalents aligned at metaphase I with a stretched-diamond arrangement similar to the wild type, indicating normal mono-orientation of sisters in the presence of chiasmata. Mono-orientation at metaphase I is confirmed by the observation of normal meiosis II, with five chromatid pairs aligned at metaphase II plates (Figure 5, Figure S2). This shows that the presence of chiasmata promotes monopolar orientation, and that the identified mutations that promote bipolar orientation of univalents do not weaken monopolar orientation enough to provoke bipolar orientation of sister kinetochores in a bivalent context.

**Figure 5.**
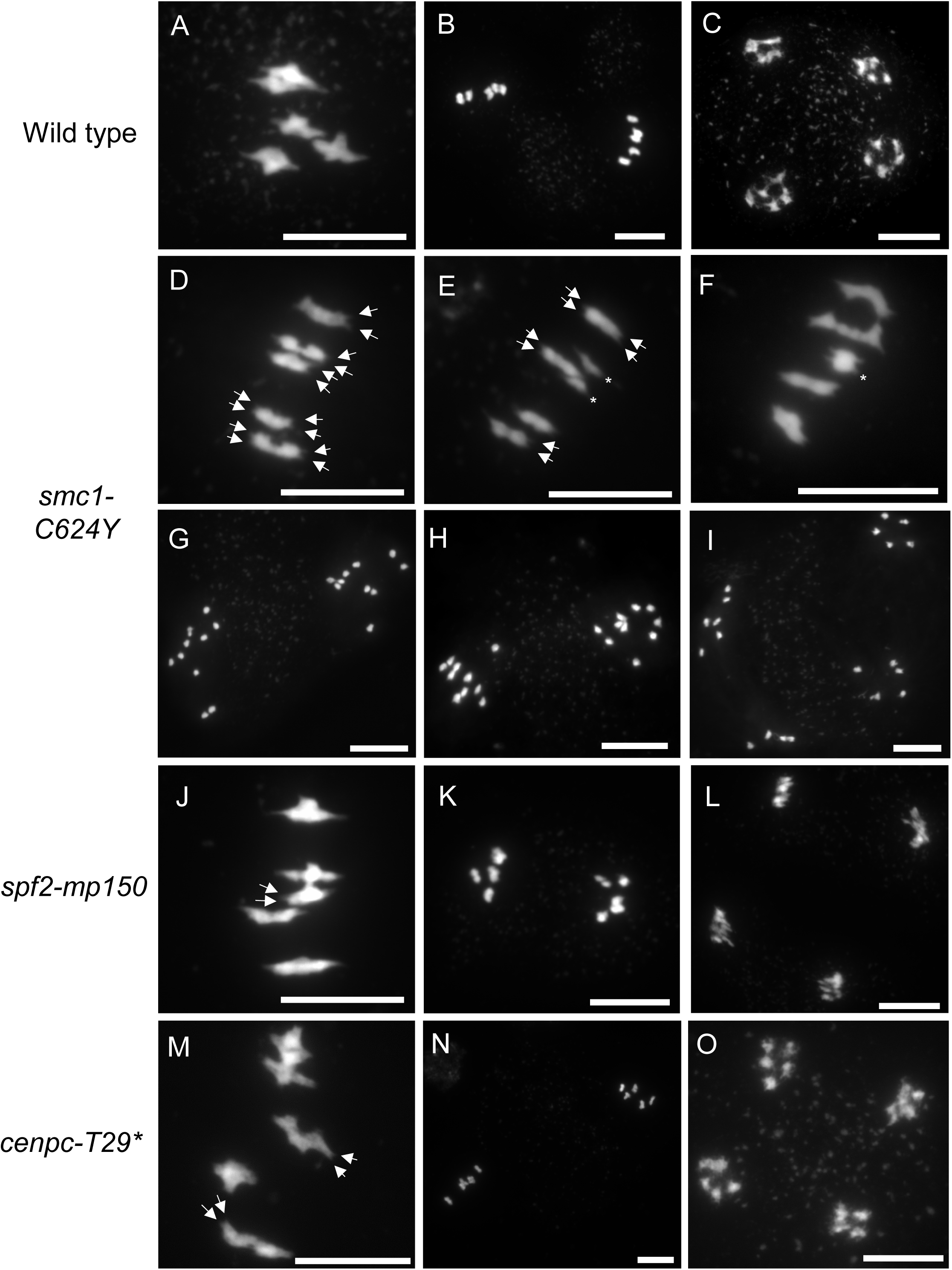
Chromosome spreads of single mutant male meiocytes. (A) Wild type metaphase I. (B) wild type metaphase II (C) wild type telophase II. (D) *smc1-C624Y* metaphase I (arrows indicate centromere splitting). (E) *smc1-C624Y* with two univalents (indicated by asterisks; arrows indicate centromere splitting). (F) *smc1-C624Y* metaphase I. (G) *smc1-C624Y* metaphase II with complete cohesion loss following meiosis I (left: 10 chromatids, right: 10 chromatids). (H) *smc1-C624Y* metaphase II with incomplete cohesion loss following meiosis I (left: 8 chromatids and 1 univalent, right: 10 chromatids). (I) *smc1-C624Y* telophase II. (J) *spf2-mp150* metaphase I with five bivalents (arrows indicate centromere splitting). (J) *spf2-mp150* metaphase I. (K) *spf2-mp150* metaphase II. (L) *spf2-mp150* telophase II. (M) *cenpc-T29\** metaphase I (arrows indicate centromere splitting). (N) *cenpc-T29\** metaphase II. (O) *cenpc-T29\** telophase II. Scale bar: 10µm.

The *smc1-C624Y* single mutant showed strongly reduced fertility and meiotic defects (Figure 2E and Figure 5G-L). This is in contrast with null smc1 mutants, which are not viable ^32^. In most *smc1-C624Y* metaphase I cells, five bivalents aligned with an arrangement similar to the wild type, suggesting efficient DNA double-strand breaks and crossover formation, and normal mono-orientation of sisters in the presence of chiasmata. However, pairs of univalents were infrequently observed (one pair of univalents in 4/43 cells), suggesting an incomplete implementation of crossovers (Figure 5H, asterisks). These univalents aligned at metaphase I in a bipolar manner, consistent with a defect in maintaining monopolar orientation of univalents. Bivalents, including in the same cells, showed a shape similar to the wild type, strongly suggesting a maintained monopolar orientation. This confirms that chiasmata themselves promote bipolar orientation, and not upstream events such as the presence of SPO11 or the formation of double-strand breaks. In rare occasions, bivalents adopt a shape suggestive of bipolar orientation of pairs of sisters despite the presence of chiasmata (asterisk in figure 5F) (see also below).

### Protection of centromeric cohesion is impaired in *smc1-C624Y* and in *sgo1 sgo2*

At metaphase II, *smc1-C624Y* meiocytes showed separated sister chromatids, with up to ten free chromatids on each plate (Figure 5 G-H), rather than five aligned chromatid pairs as in wild-type (Figure 5B). This reveals a severe premature loss of sister chromatid cohesion in *smc1-C624Y*. This suggests a failure of protection of peri-centromeric sister chromatid cohesion from Separase at anaphase I, which is required for the second division in wild-type. The same defect is observed in *sgo1 sgo2* ^8^. This shows that in *smc1-C624Y* and in *sgo1 sgo2*, both monopolar orientation and protection of centromeric cohesion are affected.

### Mutations in *SMC1, SPF2, and DCC1* cause the splitting of sister kinetochores at metaphase I

As mentioned above, in chromosome spreads of metaphase I cells, DAPI-stained bivalents resemble stretched diamonds in both wild-type and mutants. In wild-type centromeres observable at the edges of bivalents appear as a single sharp tip, suggesting a close association of sister centromeres (Figure 6). In contrast, 53% of bivalents edges in *smc1-C624Y* appear split into two tips at metaphase I. The same observation was made for in *dcc1* and in *spf2*, through at lower frequencies (Figure 6). In *cenpc-T29\**, separated centromeres are only observed in 2% of centromere tip pairs, comparable to the wild type and in clear contrast to *dcc1* or *smc1-C624Y* (Figure 6B). To test whether separation of DAPI signals at the tips of bivalents represented separation of kinetochores, we performed immunostaining of the inner kinetochore component on bivalents, using a protocol that maintains the cellular 3D organization ^46^. In wild type, the kinetochore pairs appear as adjacent signals, with an average distance between the two signal pics of 340nm. In *smc1-C624Y* the pairs of CENH3 signal appear more separated, with an average distance of 500nm (Figure 7 A-B). This confirmed the splitting of sister kinetochores at metaphase I, despite monopolar orientation. Immunostaining of CENH3 in *dcc1-1* and *spf2-mp150* also revealed a larger separation than in wild-type, with an average 460nm and 410nm respectively between sister CENH3 signals (Figure 7), similar to *smc1-C624Y* but at a reduced average distance of separation. The observed separation of sister kinetochores prior to the first division in *smc1-C624Y* combined with its complete loss of cohesion after the first division in *smc1-C624Y* (Figure 5H) suggests that sister chromatid cohesion is weakened in *smc1-C624Y*, notably in the vicinity of centromeres, which in turn could affect the robustness of monopolar orientation. DCC1 and SPF2 may promote monopolar orientation in a similar manner, given that the milder kinetochore separation in their mutants compared to *smc1-C624Y* and a concordant lower effect on monopolar orientation (Figure 2-4).

**Figure 6.**
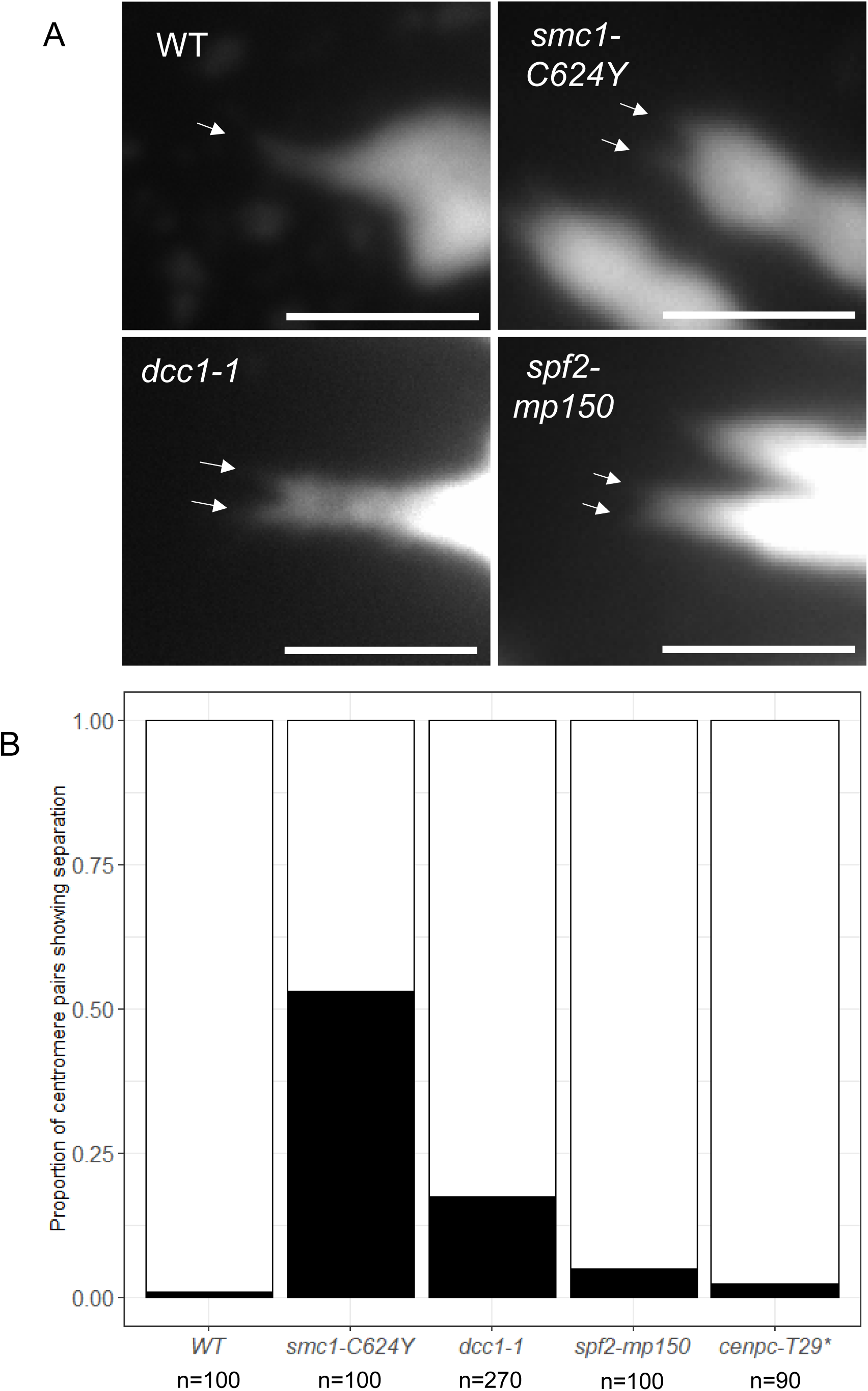
Centromere splitting in single mutant male meiocytes. (A, top left) Zoomed detail of centromeric region of wild type bivalents at metaphase I and. (A, top right) zoomed detail on *smc1-C624Y* bivalent at centromere region; arrows indicate two sister centromeres. (A, bottom left) zoomed detail of *spf2-mp150* bivalent at centromere region; arrows indicate two sister centromeres. (A, bottom right) zoomed detail of *dcc1-1* bivalent at centromere region; arrows indicate two sister centromeres. (B) Quantification of split centromeric regions in wild type and monopolar orientation mutants; each data point represents one centromeric region. Scale bar: 2.5 µm.

**Figure 7.**
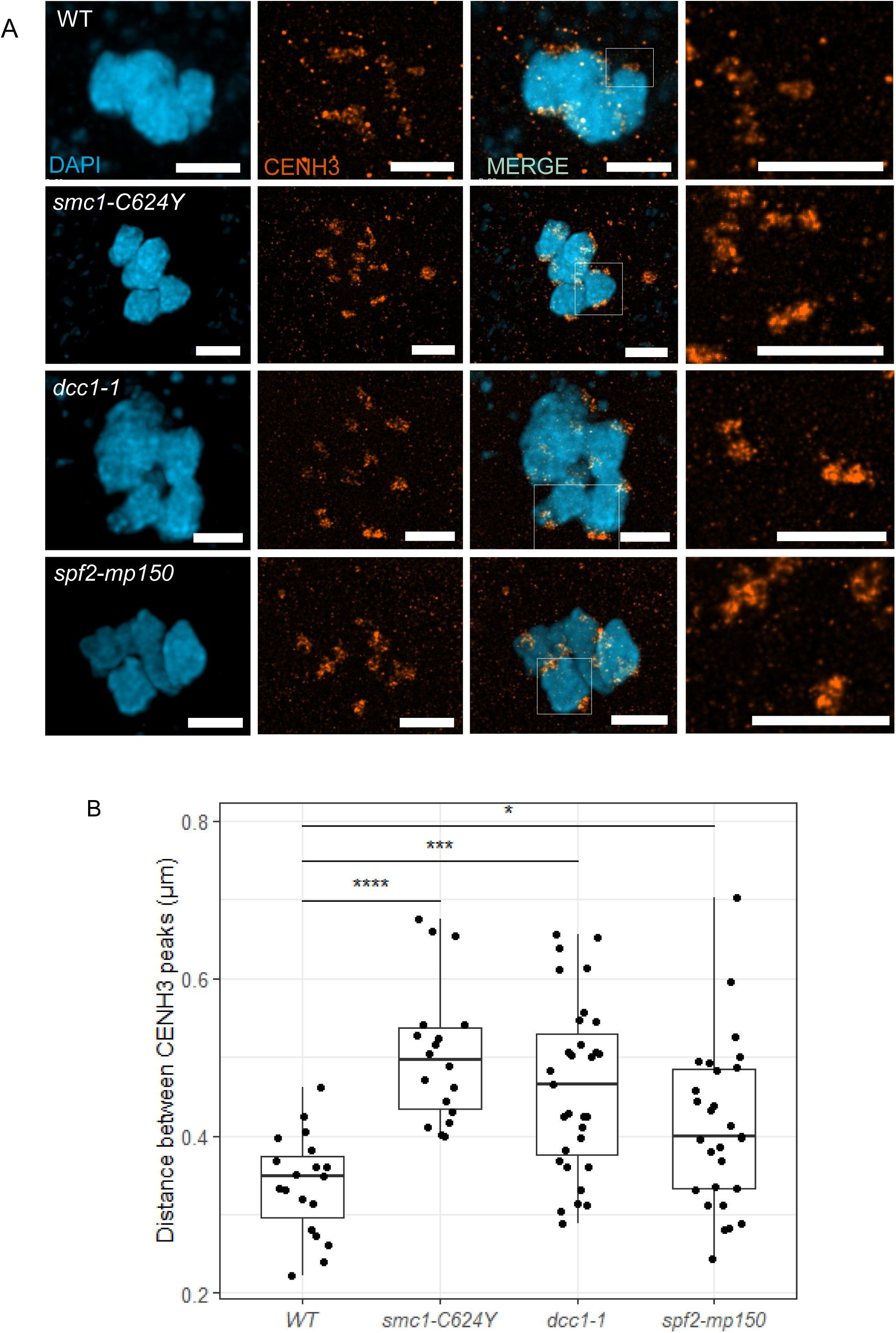
Inter-sister kinetochore distance in co-orientation mutants at metaphase I. (A) Confocal/STED images of metaphase I bivalents in the wildtype and monopolar orientation mutants; blue = confocal DAPI, orange = confocal+STED CENH3. (B) Quantification of the distance between STED CENH3 signals in the wildtype and monopolar orientation mutants, where one data point represents one pair of CENH3 signals. Scale bar: 2 µm.

### Reduced levels of cohesin on metaphase I bivalents in *dcc1* and *spf2*

To investigate whether meiotic Cohesin is affected in the described mutant alleles prior to the first meiotic division, we performed immunostaining of the meiosis-specific cohesin subunit REC8 on metaphase I bivalents. Bivalents in *dcc1-1* show a significant reduction of REC8 cohesin both at centromeres and broadly on chromosomes relative to the wild type, which is consistent with the described role of DCC1/CTF18 in facilitating cohesion establishment during replication (Figure 8 C, D). This suggests that cohesion establishment factors may promote the robustness of monopolar orientation-relevant cohesin pools by ensuring sufficient cohesin is loaded during replication. Bivalents in *spf2-mp150* similarly showed a significant reduction in meiotic cohesin broadly on chromosomes and at centromeres in *spf2* (Figure 8). This implicates the deSUMOylase SPF2 in cohesion protection, which in turn would support close association of kinetochores and monopolar orientation. REC8 levels on *cenpc-T29\** bivalents showed a slight reduction relative to the wildtype (Figure 8), suggesting that CENP-C also promotes cohesin levels at meiosis I.

**Figure 8.**
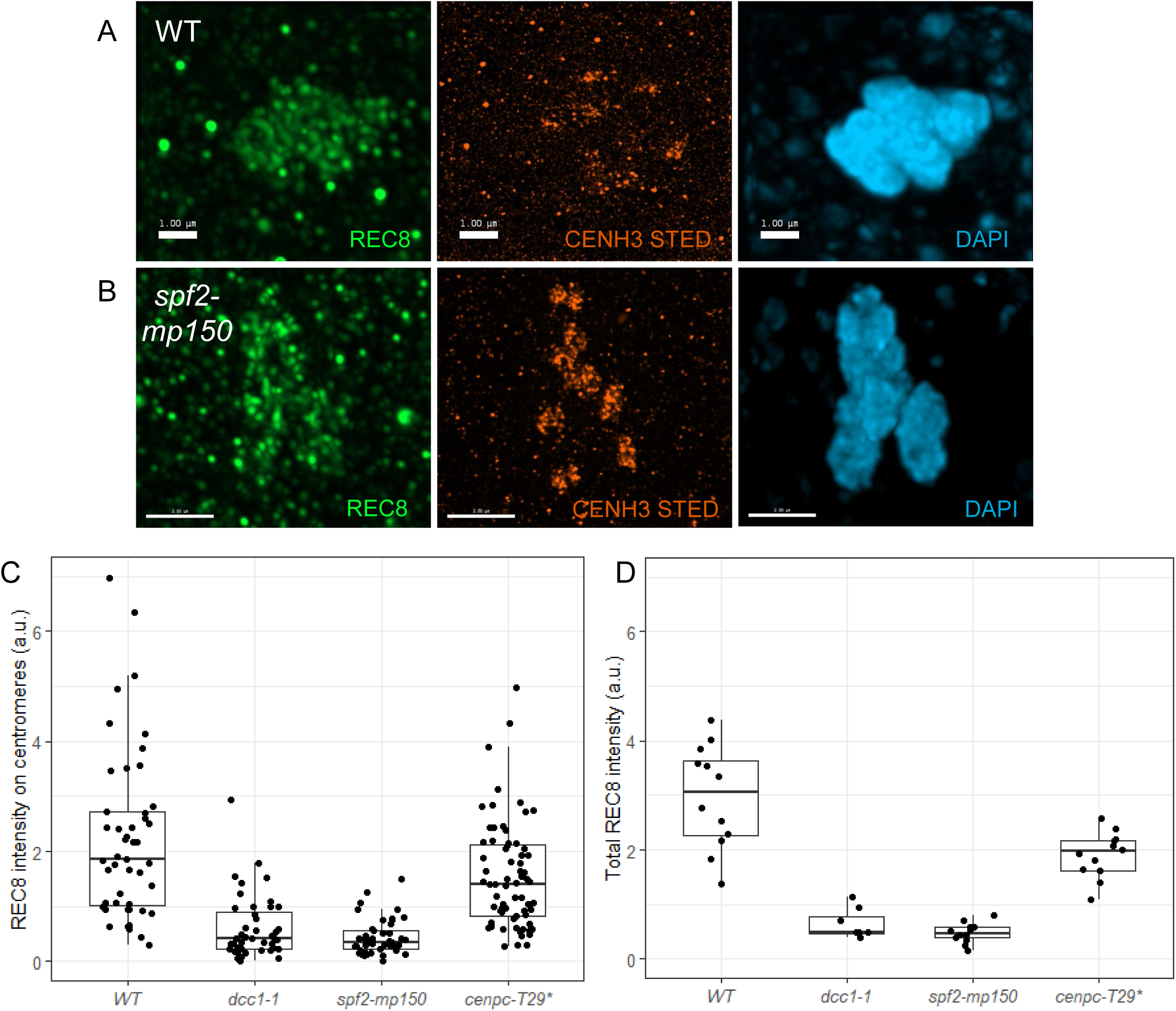
Meiotic cohesin at centromeres and chromatin in monopolar orientation mutants. (A) Confocal/STED images of wildtype metaphase I bivalents; green = confocal REC8, orange = STED CENH3, blue = confocal DAPI. (B) Confocal/STED images of *asp2-mp* metaphase I bivalents. (C) Quantification of normalized REC8 signals at centromeric regions; each data point represents one centromeric region. (D) Quantification of normalized REC8 signals on chromatin; each data point represents one metaphase I cell.

### Epistasis analysis supports a cohesion-dependent mechanism of monopolar orientation

We next combined monopolar orientation-affected alleles and assessed their meiotic behavior. The *sgo2* mutant does not show any visible meiotic defect on its own, but enhances the cohesion defects of *sgo1*, provoking a complete loss of sister chromatid cohesion after anaphase I ^8^. Strikingly, combining *sgo2* with *cenpc-T29\** also provoked a severe loss of sister chromatid cohesion at telophase I (Figure 9C), while both single mutants do not show this defect (Figure 5M-O). This implicates that CENP-C promotes the maintenance of centromeric sister chromatid cohesion during meiosis. Similarly, the *spf2-4 cenpc-T29\** double mutant showed loss of sister chromatid cohesion after anaphase I (Figure 9F), while none of the single mutant show this defect. These epistatic interactions implicate both *SPF2* and *CENP-C* in centromeric cohesion protection. Together, the findings implicate both CENP-C and SPF2, together with SGOs, in monopolar orientation and in the maintenance of centromeric cohesion until metaphase II.

**Figure 9.**
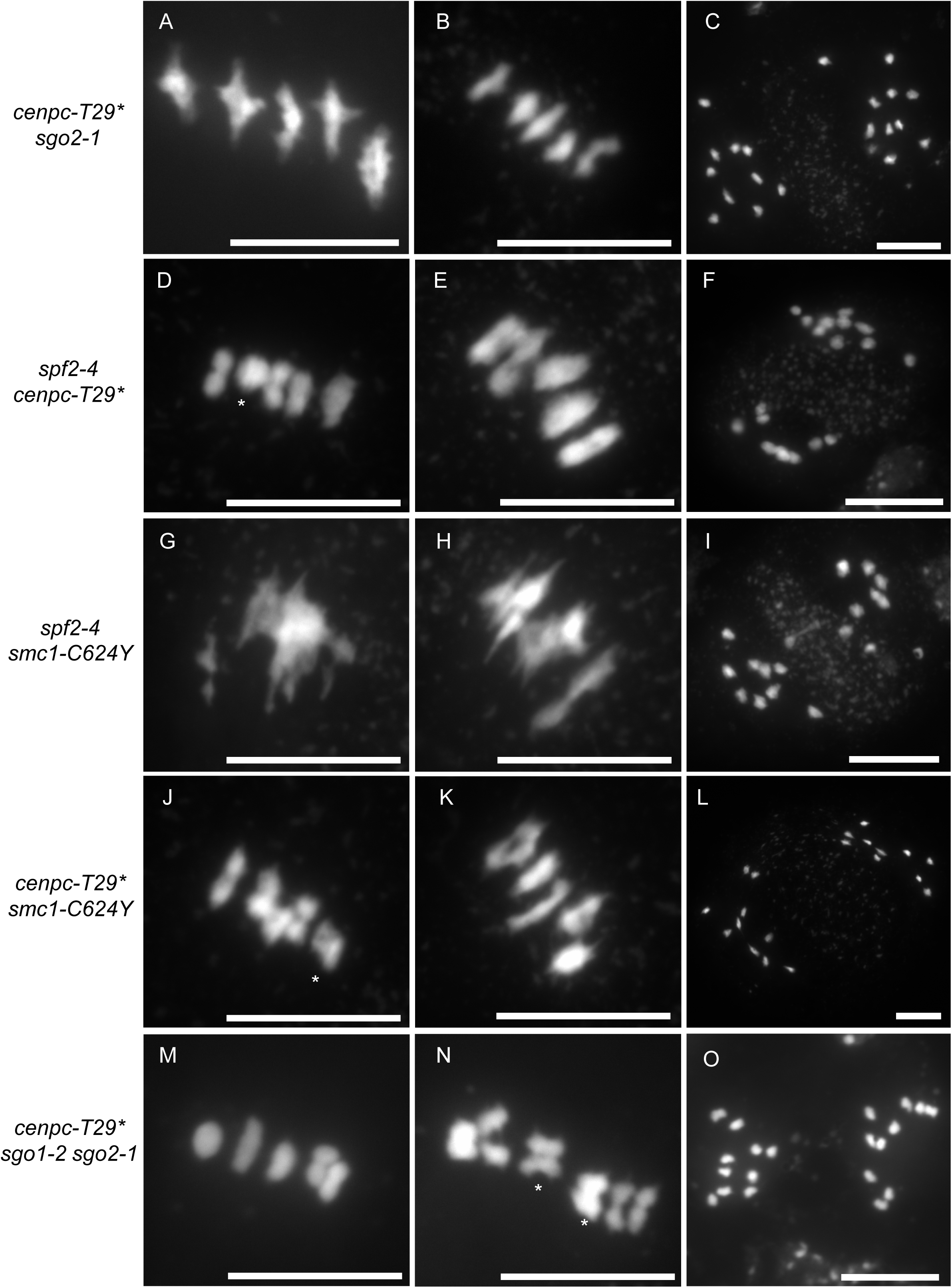
Chromosome spreads of monopolar orientation double mutants. (A-C) *cenpc-T29* sgo2-1*. (D-F) *cenpc-T29* spf2-4*. (G-I) *spf2-4 smc1-C624Y*. (J-L) *cenpc-T29* smc1-C624Y.* (M-O) *cenpc-T29* sgo1-2 sgo2-1*. (A,D,G,J,M) metaphase I. (B,E,H,K,N) metaphase II. (C,F,I,L,O) telophase II. Scale bar: 10 µm.

*The spf2-4 smc1-C624Y* double mutant displays a stretched and aberrant bivalent structure at metaphase I (Figure 9G), reminiscent of mutants with DNA double strand-break repair defects, notably null *rec8* ^11^. This suggests that in this double mutant Cohesins are decreased to a level that affects the recombination process. Loss of sister chromatid cohesion at metaphase II is also observed in *spf2-4 smc1-C624Y* and in *cenpc-T29* smc1-C624Y* (Figure 9I and 9L), like in the single *smc1-C624Y*. Similarly, complete loss of sister chromatid cohesion at metaphase II is observed in *cenpc-T29* sgo1 sgo2*, like in *sgo1 sgo2* (Figure 9O) ^8^.

At metaphase I, distinct shapes were observed for bivalents in *cenpc-T29* sgo1-2 sgo2-1*, *cenpc-T29* smc1-C624Y* and *cenpc-T29* spf2-3* double mutants (Figure 9M-N,J-K,D-E). The shape of these bivalents diverges from that of the wildtype typical stretched-diamond-shape. They appear instead as rounded oval or squares. Some bivalents with a square shape are strongly suggestive of a bipolar orientation of the two pairs of sister chromatids (Figure 9D, 9J, 9N asterisks). This is reminiscent of the shape of bivalents with bi-orientation in human oocytes^47^. Thus, in *cenpc-T29* sgo1 sgo2*, *cenpc-T29* smc1-C624Y* and *cenpc-T29* spf2* mutants, monopolar orientation appears to be sufficiently affected to allow bi-orientation of chromatid pairs within a bivalent, overcoming the pro-monopolar effect of chiasmata. This is also consistent with the complete loss of cohesion observed at metaphase II in these three double mutants. Taken together, this epistasis analysis implicates both SPF2 and CENP-C in sister chromatid cohesion and its maintenance until meiosis II. This supports a model in which SPF2 and CENP-C - together with Cohesins and their regulators DCC1/CTF18 and SGOs-promote the monopolar orientation of kinetochores at meiosis in a cohesion-dependent manner.

## Discussion

### Monopolar orientation is highly dependent on Cohesins

Monopolar orientation in yeasts and metazoans appears to be largely promoted by the specific regulation of meiotic cohesin in the vicinity of centromeres ^10,16,18^. We demonstrate here, that in addition to the meiotic-specific cohesin subunit REC8, the other cohesin subunits *SMC1*, *SMC3,* and *SCC3* promote monopolar orientation in Arabidopsis. These three subunits are each encoded by a single gene and are essential for mitosis, but we identified mutations in each of them that weaken monopolar orientation, yet remain viable. We further find that CTF18 and DCC1 support monopolar orientation. CTF18 and DCC1 belong to the replication machinery and play a role in establishing sister chromatid cohesion. Accordingly, we observed a reduced level of cohesins on *dcc1* meiotic chromosomes. Arabidopsis *ctf18* and *dcc1* loss-of-function mutations are viable and fertile, but promote monopolar orientation in univalents. Altogether, this suggests that monopolar orientation requires a level of cohesion fidelity beyond that of other functions of Cohesins at mitosis and meiosis.

We show that the two plant Shugoshins SGO1 and SGO2 support monopolar orientation. Shugoshin is a conserved family of proteins that protect cohesins during meiosis by shuttling protein phosphatase 2A (PP2A) to pericentromeres to dephosphorylate REC8 and prevent its cleavage by Separase ^6,48,49^. This protection of cohesin in the centromeric region is essential for alignment and balanced distribution of chromatids at the second division. Both Arabidopsis SGO1 and SGO2 function in centromeric cohesion protection during meiosis, in a partially redundant manner ^8^. Mutation of *SGO1* leads to a partial premature loss of centromeric cohesion, mutation of *SGO2* does not, and a complete premature loss of cohesion is observed in the double mutant. Similarly, we show here that both SGO1 and SGO2 promote monopolar orientation, in a partially redundant manner. However, their relative importance appears to be reversed, with a higher frequency of bipolar orientation of univalents at metaphase I in *sgo2* than in *sgo1* and a higher frequency when both are disrupted. The observation that SGO1 and SGO2 have opposite relative importance for monopolar orientation and cohesin protection suggests that these two proteins have distinct targets. One attractive model is that SGO1 and SGO2 protects two distinct pools of cohesins, one promoting monopolar orientation at meiosis I and the other ensuring centromeric cohesion at meiosis II.

### AtCENP-C promotes mono-orientation and cohesion protection

The inner kinetochore protein CENP-C is a crucial scaffold component that regulates kinetochore assembly during mitosis and meiosis ^50^. We show here that CENP-C also has a role in imposing mono-orientation during meiosis. The *cenpc-T29\** allele is able to promote equational segregation of univalents at anaphase I. CENP-C appears to also promote cohesion, as REC8 level are reduced on *cenpc-T29\** metaphase I chromosomes, and combining the *cenpc-T29\** with either *sgo2* or *spf2* triggers the early loss of sister chromatid cohesion after the first division. Taken together, CENP-C likely promotes monopolar orientation during meiosis I in a cohesion-dependent manner.

### deSUMOylation promotes monopolar orientation

Our findings implicate SPF2, one of the Arabidopsis homologues of the yeast Ulp2, in promoting monopolar orientation. Reduced levels of Cohesin and splitting of kinetochores at metaphase I, together with premature loss of cohesion in *spf2 cenpc-T29\**, suggest that SPF2 also participates in cohesin protection, and could also promotes monopolar orientation in a cohesion-dependent manner. SUMOylation is known to have diverse roles in coordinating chromosome segregation in both mitosis and meiosis through means of kinetochore assembly, regulation of error correction, and cohesin establishment and maintenance ^51–55^. Ulp2 has been shown to prevent the build-up of SUMO chains on SMC complexes including cohesin in budding yeast ^56^. Loss of SPF2 may lead to premature turnover of Cohesin, weakening both centromeric cohesion and monopolar orientation. As SUMOylation is pervasive at meiosis, one cannot exclude cohesin-independent roles of SPF2 at kinetochores ^57^. Notably, Ulp2 also prevents SUMOylation-mediated turnover of Condensins, which are also promote monopolar orientation in *S. Cerevisiae* ^56,58^.

### Split kinetochores, chiasmata and monopolar orientation

All mutations that we describe here (*smc1, smc3, scc3, ctf18, dcc1, ctf18, sgo2, sgo1, cenpc, spf2*) were able to suppress monopolar orientation of sister kinetochores of achiasmatic univalents (*spo11* metaphase I). However, in all of them, monopolar orientation was restored by the presence of chiasmata (bivalents), strongly supporting the role of chiasmata in promoting monopolar orientation ^59^. Although sister kinetochores were oriented to the same pole in bivalents, like in wild-type, they were more separated from each other than in the wild-type. This was also associated with reduced cohesin levels. This suggests a model in which all these mutations reduce Cohesin levels, loosening the normally tight association between the sister-kinetochores. This loose association is still sufficient to promote monopolar orientation in the presence of chiasmata, but not in univalents where only bipolar orientation can generate the tension needed to stabilize metaphase spindle. This is consistent with observations in mouse oocytes that demonstrated that specific extinction of REC8 in the immediate vicinity of centromeres causes a significant increase in inter-sister kinetochore distance in metaphase I bivalents; this separation alone is not sufficient to reversion mono-orientation in bivalents, but effectively does so when univalents are present at the metaphase I plate ^18^. Combining mutations, and presumably further destabilizing Cohesins led to a range of meiotic defects. In *spf2 smc1-C624Y*, we observed aberrant chromosomal structures, similar to that observed in *rec8* which are caused by unrepaired recombination intermediates ^11,60^. This highlights the essential role of cohesin in homologous recombination. In several combinations, we also observed premature loss of cohesion before metaphase I, suggesting an insufficient maintenance of centromeric cohesin. Finally, in three mutant combinations (*cenpc-T29* sgo1 sgo2*, *cenpc-T29* smc1-C624Y* and *cenpc-T29* spf2*), we observed rotated bivalents, strongly suggesting that monopolar is sufficiently affected to allow bipolar orientation of the two sister chromatid pairs even in the presence of chiasmata within the bivalent structure.

### Are MOKIRs specific to Opisthokonta?

The findings presented here implicate a significant number of novel players in the proper mono-orientation of kinetochore complexes prior to the first meiotic division. While long recognized as an essential and unique feature of meiosis ^61^, mechanisms of monopolar orientation have remained elusive, notably in the plant kingdom. In *opisthokonta*, the eukaryotic branch that contain both fungi and animals, it appears that the meiotic specific protein MOKIRs plays a central role. MOKIRs stands for Meiosis One KInase Regulator and is represented by Moa, Spo13 and MEIKIN in *S. cerevisiae*, *S. pombe and* mammals, respectively ^16,25,62–64^. These mostly intrinsic disordered proteins are considered as orthologues despite limited sequence conservation, and act as a meiosis-specific shuttle for Polo kinase localization to the kinetochore ^13,64^. Mouse oocytes and spermatocytes lacking MEIKIN show signs of defective mono-orientation, with frequent splitting of kinetochores prior to anaphase I and bi-orienting kinetochores in cells lacking chiasmata, reminiscent of the series of mutants described here. However, none of the Arabidopsis genes we identified here correspond to the Moa/Spo13/MEIKIN portrait. We confirmed that monopolar orientation in Arabidopsis depends on the regulation of Cohesins, like in *Opisthokonta*, but did not find any meiosis-specific protein involved in the process, except the meiotic cohesin subunit REC8. Several reasons can explain how we could have missed a MOKIR plant representative, such as gene duplication of lack of mutation saturation, but this raises the possibility that it does not exist. We also failed to identify a MOKIR candidate in the plant kingdom using sensitive homology search. It should be noted that the Polo kinase/PLK is absent in plants ^65^. One possibility is that plants use a different kinase for this function, and that a meiotic-specific shuttle equivalent to MOKIR exists. One may alternatively envisage that MOKIR arose in the *Opisthokonta* clade to reinforce the meiotic chromosomal architecture, and that the cohesin REC8 might be sufficient to confer meiotic specificity in plants and possibly in the ancestral eukaryote.

## Materials and methods

### Plant materials and growth conditions

*Arabidopsis thaliana* plants were grown in greenhouses or growth chambers under 16 hour day/8 hour night conditions at 20 degrees C. In addition to the EMS and CRISPR alleles produced in this study we used the following genetic material:

*osd1-3* ^66^, *rec8-3*/SAIL_807_B08 ^28^, *spo11-1-3*/SALK_146172 ^67^, *spo11-2-3*/GABI_749C12 ^68^, *smc1-1*/SALK_017437 ^32^, *sgo1-2* and sgo2-1 ^8^, *spf2-3*/asp2-2/ SALK_140824 ^69^, *spf2-4/asp2-3/*SAIL_182_F02 ^70^, *ctf18-3/*SALK_043339 ^71^, *glabra1-hc1* ^72^, *GFP-tailswap* ^26^,*pans1-1*/Salk_070337 ^8^

### Mutagenesis

Seeds were incubated for 17 h at room temperature with gentle agitation in 5 mL of 0.1% (v/v) EMS. Neutralization was performed by adding 5 mL sodium thiosulfate 1M for 5 min. Three milliliter water was added to make the seeds sink. The supernatant was removed and the seeds were washed three times for 20 min with 15 mL of water before sowing.

### CRISPR-Cas9 mutagenesis of *AtDCC1*

To assemble a complete CRISPR-Cas9 binary vector, four guide RNAs were first designed spanning the genomic sequence of AT2G44580 using CRISPOR ^73,74^. Cloning of the guide RNAs into the final Cas9-containing binary vector was carried out using the MoClo Golden Gate cloning toolkit ^75^ and integrating an intron-containing Cas9 ^76^. Briefly, guide RNAs were synthesized as an oligonucleotide comprising the target sequence and BsaI restriction enzyme recognition sequence. In the first cloning reaction, each guide was independently fused to an AtU6 promoter while simultaneous adding complementary BpiI recognition sites. In the final cloning reaction, all four guides were assembled with a selection marker (FastRED) and Cas9. This completed construct was then transformed into A. tumefaciens, and harvested cells were used to transform Col-0 A. thaliana plants using the floral dip method ^77^. Positive transformants were selected by screening seeds for FastRED fluorescence.

### Chromosome preparations

Young inflorescences were harvested from Arabidopsis plants and immediately fixed in 3:1 ethanol:acetic acid. Flower buds roughly 0.5mm in size were isolated from inflorescences and washed in citrate buffer, then incubated in driselase enzyme mixture (Sigma) for 2 hours at 37 degrees C. Four to five digested buds were transferred to a clean slide and macerated with a bent dissection needle. Roughly 15uL of 60% acetic acid was added to the mixture and stirred gently at 45 degrees C on a hotplate for 1 minute. The slide was then flushed with ice-cold 3:1 fixative first around the droplet and then directly^78^. Slides were left to dry tilted at room temperature, then 10uL of DAPI in mounting media was applied to the slide and a coverslip was added. Chromosomes were visualized using a Zeiss Axio Observer epifluorescence microscope.

### Immunocytology

Slides that preserve the 3D structure of freshly fixed meiocytes were prepared and used for immunocytology using published methods^79^. Briefly, fresh inflorescences were collected from Arabidopsis plants, dissected to expose young athers, and fixed in paraformaldehyde for 30 minutes under vacuum. Buds roughly 0.5mm in size were selected and digested in driselase cellulase/pectinase enzyme mixture for 30 minutes to remove cell walls. Anthers were then isolated and slightly ruptured to release meiocytes, then squashed with a polyacrylamide gel layer onto coverslips. Coverslips were stained with DAPI to stage meiocytes, then treated for 48 hours at 4 degrees C with primary antibodies for CENH3 (anti-chicken, 1:500) or REC8 (anti-rabbit, 1:250) prepared in blocking buffer. Secondary antibodies were then incubated for 24 hours at 4 degrees C before addition of mounting media with DAPI and the sealing of slides with nail polish. Slides were imaged using an Abberior Facility Line STED microscope using a 2D STED depletion laser. STED images were processed using Huygens Essential software to perform deconvolution and distances between STED CENH3 signals were measured using Fiji (ImageJ) on maximum intensity projections. To quantify relative fluorescence, non-deconvolved sum projection images were used in Fiji. Regions of interest (ROIs) were drawn over confocal CENH3 signals for centromeres or DAPI signals for chromatin. Mean background confocal REC8 signal was subtracted from mean REC8 signal on centromeres or chromatin, and then normalized by dividing this value by background-corrected CENH3. Thus, (S_REC8_-S_REC8 bg_)/(S_CENH3_-S_CENH3 bg_), where S is the mean signal and bg is background.

## Acknowledgements.

We thank Dipti Vernekar for her suggestions on the manuscript and Neel Ajay Shah for technical help. This work was supported by core funding from the Max Planck Society. D.K.S. is supported by the DBT-Ramalingaswami Re-entry Fellowship. This work has benefited from the support of IJPB’s Plant Observatory platforms PO-Plants.

---

**Figure S1.**
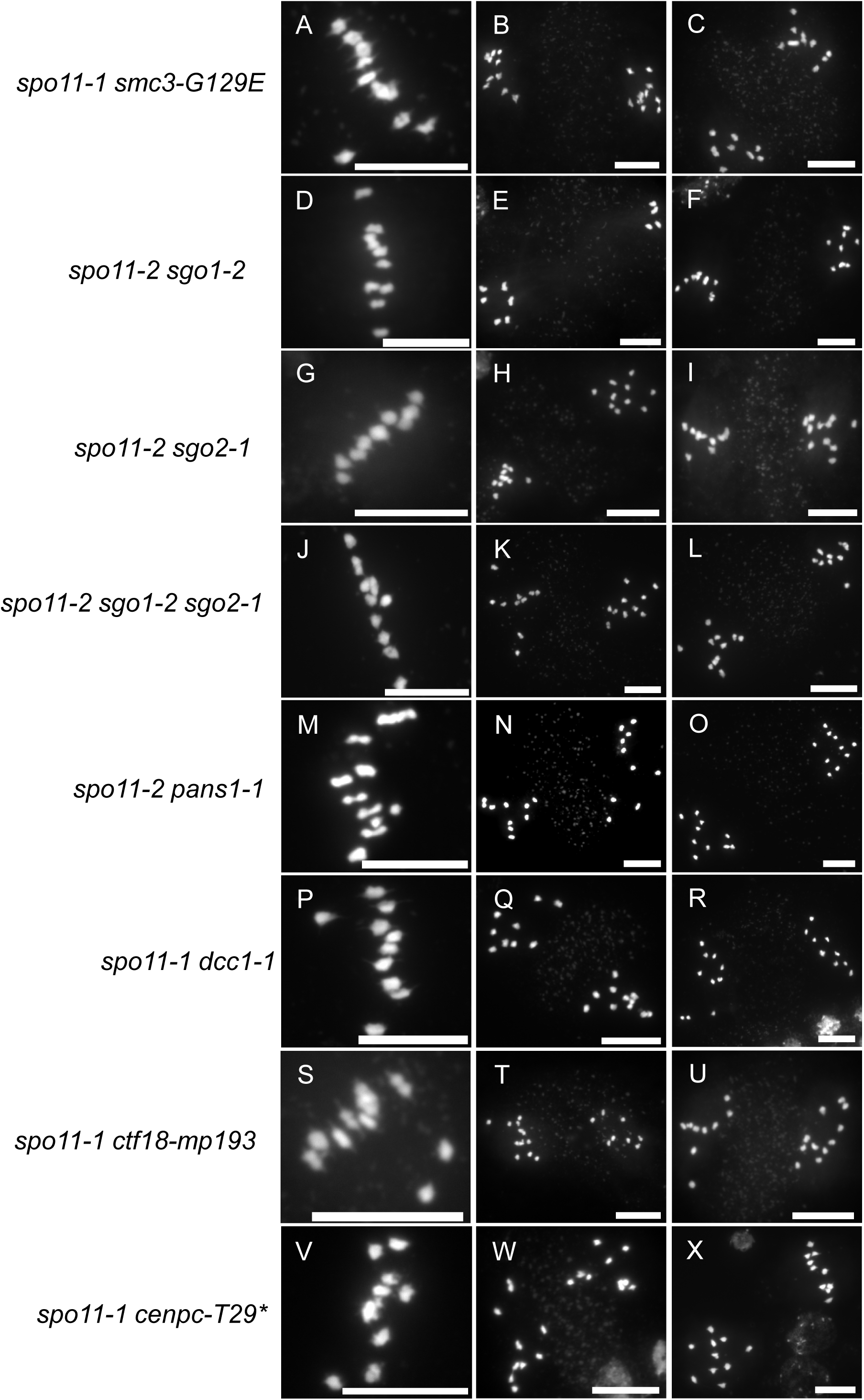
Male meiocyte chromosome spreads in monopolar mutants in the achiasmatic *spo11* background. (A) *spo11-1 smc3-G129E* metaphase I. (B-C) *spo11-1 smc3-G129E* metaphase II. (D) *spo11-2 sgo1-2* metaphase I. (E-F) *spo11-2 sgo1-2* metaphase II. (G) *spo11-2 sgo2-1* metaphase I. (H-I) *spo11-2 sgo2-1* metaphase II. (J) *spo11-2 sgo1-2 sgo2-1* metaphase I. (K-L) *spo11-2 sgo1-2 sgo2-1* metaphase II. (M) *spo11-2 pans1* metaphase I. (N-O) *spo11-2 pans1* metaphase II. (P) *spo11-1 dcc1-1* metaphase I. (Q-R) *spo11-1 dcc1-1* metaphase II. (S) *spo11-1 ctf18-mp193* metaphase I. (T-U) *spo11-1 ctf18-mp193* metaphase II. (V) *spo11-1 cenpc-T29\** metaphase I. (W-X) *spo11-1 cenpc-T29\** metaphase II. Scale bar: 10µm.

**Figure S2.**
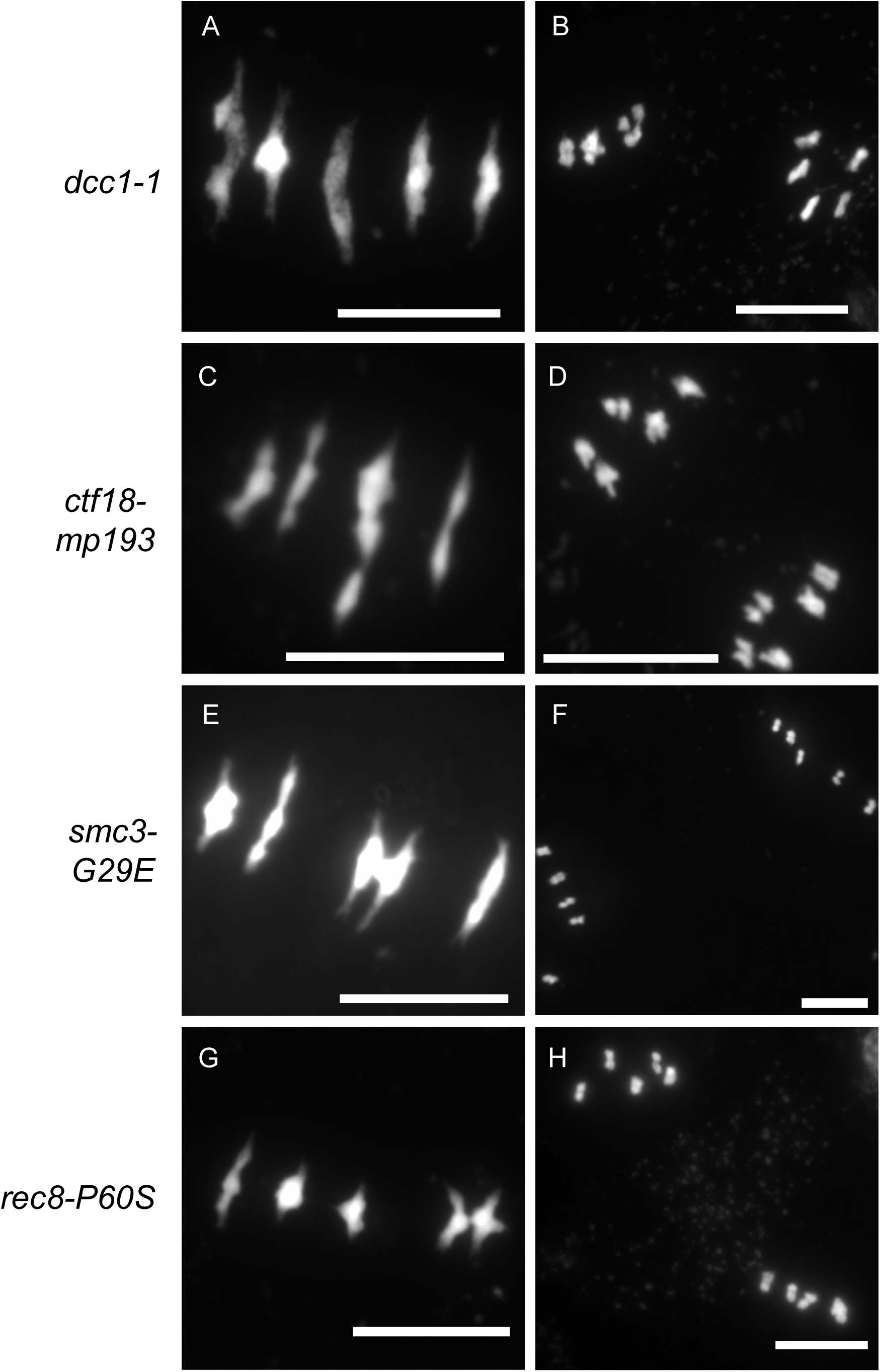
Male meiocyte chromosome spreads in single monopolar mutants. (A) *dcc1-1* metaphase I. (B) *dcc1-1* metaphase II. (C) *ctf18-mp193* metaphase I. (D) *ctf18-mp193* metaphase II. (E) *smc3-G29E* metaphase I. (F) *smc3-G29E* metaphase II. (G) *rec8-P60S* metaphase I. (H) *rec8-P60S* metaphase II. Scale bar: 10µm.

